# PMScanR: an R package for the large-scale identification, analysis, and visualization of protein motifs

**DOI:** 10.1101/2025.05.23.655703

**Authors:** Jan Pawel Jastrzebski, Monika Gawronska, Wiktor Babis, Miriana Quaranta, Damian Czopek

**Affiliations:** Faculty of Biology and Biotechnology, University of Warmia and Mazury in Olsztyn, Olsztyn, Warmian-Masurian Voivodeship, Poland; Ecology and Genetics, Faculty of Science, University of Oulu, Oulu, Northern Ostrobothnia, Finland; Department of Biochemical Sciences “A. Rossi Fanelli”, Sapienza Università di Roma, Rome, Lazio, Italy

## Abstract

Proteins play a crucial role in biological processes, with their functions closely related to structure. Protein functions are often associated with the presence of specific motifs, which are short amino acid sequences with a specific amino acid pattern. Most bioinformatics tools focus on identifying known motifs and they lack the ability to analyze the impact of single substitutions on entire domains or motifs. To address this, we developed PMScanR, an R-based library that automates the prediction and evaluation of the impact of single amino acid substitutions on the occurrence of protein motifs in large datasets. In addition, existing tools do not support comparative analysis of multiple motifs across multiple sequences - a key feature that PMScanR was designed to provide. The package integrates various methods to facilitate motif identification, characterization, and visualization. It includes functions for running PS-Scan, a PROSITE database tool. Additionally, PMScanR supports format conversion to GFF, enhancing downstream analyses such as graphical representation and database integration. The library offers multiple visualization tools, including heatmaps, sequence logos, and pie charts, enabling a deeper understanding of motif distribution and conservation. Through its integration with PROSITE, PMScanR provides access to up-to-date motif data, making it a valuable tool for biological and biomedical research, particularly in protein function annotation and therapeutic target identification.

The PMScanR library is freely available (GPL license) on Bioconductor. The source code, precompiled library, installation instructions, tutorial files, and complete documentation can be found on GitHub: github.com/prodakt/PMScanR.

## Introduction

Proteins are key macromolecules with many biological functions, such as catalysis of enzymatic reactions, molecular transport, cell signalling and many more. Protein functions are closely related to their three-dimensional structure, which is determined by the specific amino acid sequences. Distinct functions are often associated with the presence of protein motifs - short sequences comprising a specific pattern of repetition of amino acid residues [1]. They are crucial for understanding the molecular mechanisms underlying many protein functions. For example, specific motifs enable the protein to be involved in interactions or to undergo post-translational modifications [2]. Moreover, protein motif analysis provides information on protein evolution, classification and functional annotation, particularly important when studying proteins of unknown function or in the case of sequence modifications that could potentially modify a molecule’s function. Protein motifs can also serve as disease biomarkers, helping to identify drug targets, making their study crucial in both basic and applied biomedical research. Commonly used bioinformatics tools offer limited capabilities for protein motif extraction, analysis and visualization [3]. Tools and databases such as Pfam [4], PROSITE [5] or InterPro [6] focus mainly on motif identification, and comparative analyses between multiple sequences must be performed in multi-step summaries, often as hand-prepared tables by the researcher. This manual approach process limits scalability and reproducibility, making it difficult to analyze large datasets efficiently. As a response to this challenge we have developed a library in the R environment and compiled a user-friendly library to enable analysis of protein motifs, including their identification, characterization and comparative visualization. A key goal of the tools we developed was to facilitate and automate the analysis of the effect of amino acid sequence changes (including even single substitutions) on the occurrence of protein motifs. In addition, this library allows the preparation of summary tables and visualization of large-scale comparative analysis, and the evaluation of the stability or conservativeness of motifs.

## Design and Implementation

PMScanR (Protein Motifs ScaneR) is a library designed in the R environment that integrates various amino acid sequence analysis methods using both popular biological data formats (i.e.: FASTA, GFF, ALN) and freeware popular databases and analysis programs (e.g. PROSITE with ps scan, pfam, UniProt). It works (and can be interfaced) with other Bioconductor [7] packages making it easy (by the construction of pipelines) for further analysis and data transfer to any other (analysis or visualization) tool in the R environment. In order to facilitate the use of the library, a graphical interface was developed using the shiny library in R [8], which is launched with the runPMScanRShiny() function. The developed graphical interface enables a comprehensive analysis from start to end - ranging from the identification of motifs using ps scan up to the final visualisations. Furthermore, it is possible to load previously obtained results into the interface and perform the visualisation itself. The structure of the functions implemented in the graphical interface is compatible with the functions of the library from the level of command console or R script. This gives the possibility to carry out analysis both from the command level and the graphical interface with a smooth possibility of combining these forms. Detailed documentation and usage examples are available on the package’s website: https://github.com/prodakt/PMScanR. The PMScanR package has been tested on three types of operating systems - Windows (five versions), Mac OS X (three versions), Linux (four distributions). To install PMScanR from GitHub, you can use the install github (“prodakt/PMScanR”) function from the devtools package [9]. The package includes (provides) integration with the ps scan tool, which is part of the PROSITE database. This makes it possible to identify motifs directly inside the R environment, simplifying the process and guaranteeing compatibility of the resulting file formats. The PMScanR package provides a unified runPsScan() function to perform scanning on all main operating systems, including Windows, Linux and macOS. The function automatically detects the user’s operating system if it is not explicitly specified. In addition, runPsScan() verifies the presence of necessary execute files, databases and support files by downloading them from the PROSITE server if they are not available locally. If necessary, users can also specify custom file locations. The function supports the extensive configuration options available in ps scan, enabling flexible analysis, multiple output formats and integration with Pfam data. The motif patterns are taken from the “prosite.dat” file, so the user can keep updating his own pattern base. In the simplest configuration, the user only needs to indicate sequences file in fasta format, and all further analysis will be performed without any other input parameters. The data format adopted within the analytic environment is identical between all operating systems. The user can indicate the format of the output file, but in a further step the import of all types of output formats ps scan. Functions readPM.prosite() and readPM.psa() convert all types to tabular GFF. This is the most flexible format better suited for further analyses, such as chart generation or integration with other databases. The results of the various stages of analysis are saved to text files (easy to read). The matrix of the occurrence (made by using the gff2matrix() function) of all motifs in all sequences (or only in a part of the selected ones) contains 0/1 digits so even for very large-scale analyses it should not take up much disk space and is easy to visualize and analyze in any external program in which tables can be imported as calculation sheets. The package also offers a range of functions for data visualisation to help interpret the results. These visualizations include heatmap generation, in which the input file is a matrix of motif occurrences, and then a heatmap is generated showing the distribution of motifs in the analyzed sequences. In the current version of the package, two functions have been developed for heatmap generation: matrix2hm(), which provides the ability to adjust the plot size - this facilitates comparative analysis (Fig. 1A), and the matrix2hm 2() function, which makes it easier to identify individual variations. By using the plotly library ([10]), the plots are interactive, allowing both extensive manipulation of the plot area, but also precise identification of individual results. The next type of available type of visualization is the logo generation based on the frequency of occurrence of individual amino acid residues in a selected part of the sequence (Fig. 1B), that is, the consensus of the motif sequence (prepare segments()) and the defined region in selected sequences (extract segments() and extract protein motifs()). There is also a visualization that generates a pie chart of the frequency of each motif type from the GFF format file containing information about the motif names and their locations. The pie chart shows the percentage of each motif/protein motif type in the analyzed dataset. This function uses the ggplot2 library [11]. The pie chart allows a quick assessment of motif frequencies.

**Fig 1.**
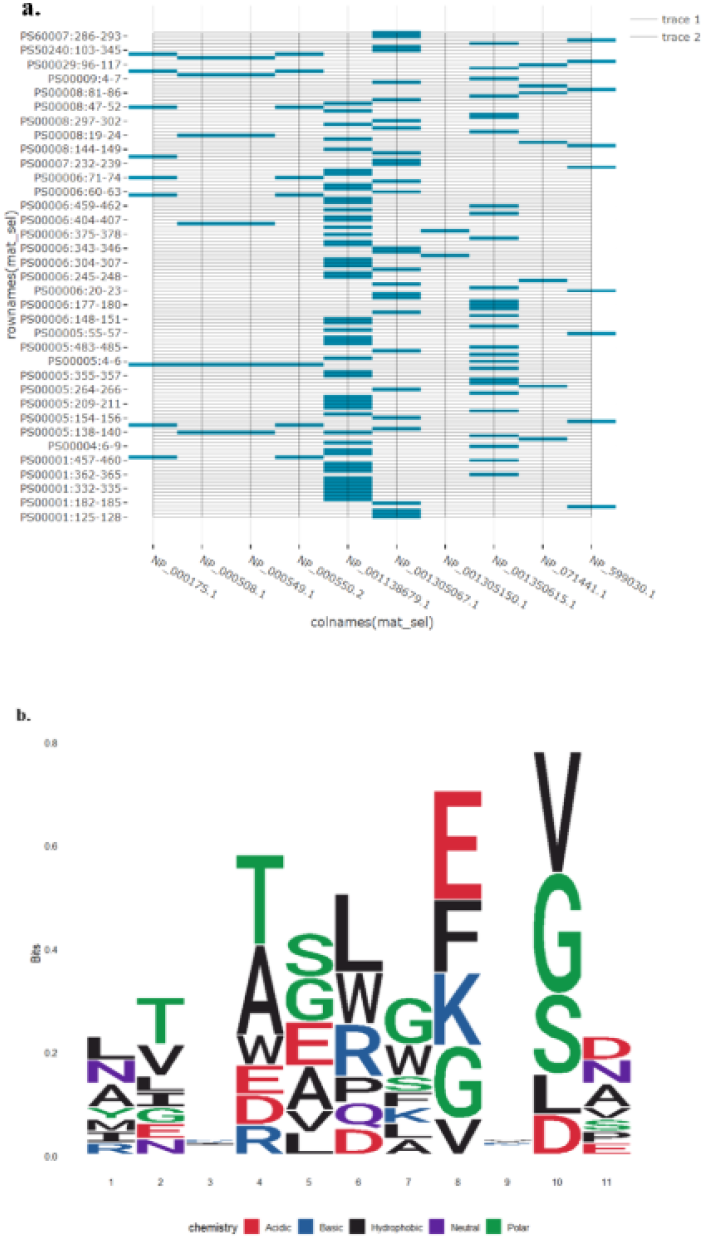
Heatmap and sequence logo of protein motifs. a. Heatmap showing the comparison of protein motifs in relation to the positions on which they occur. b. Seqlogo showing graphical representation of the sequence conservation of amino acids presents a consensus sequence and a diversity of sequences.

## Results

PMScanR was developed as a protein motif analysis package to enable easy and user-friendly analyses. The package is based on the R language, which provides flexibility and access to a wide range of analysis functions, as well as interaction with the PS-Scan tool (which is part of the PROSITE database), ggplot2, spreadsheets programs. It enables rapid and comprehensive analysis of large scale sequence data for all protein motifs deposited in PROSITE at once. PMScanR is designed for scientists and bioinformaticians who need an easy-to-use automated approach to protein motif analysis. The main functionality of the package is to enable comprehensive comparative analysis of a large number of sequences and numerous motifs simultaneously. This provides the ability to identify the potential impact of even subtle changes on protein functionality. In addition, it allows one to determine the stability and conservativeness of motifs in the compared set of sequences.

## Availability and Future Directions

The PMScanR tool is available on two open-access platforms: Bioconductor and GitHub (https://github.com/prodakt/PMScanR). On GitHub the most recent version of all source code files can be found, along with a history of all previous versions of the software. Also the tutorial files, sample input and output files, as well as several user guide files are available. The tool will continue to be developed, and future updates will focus on increasing functionality toward 3D model analysis, new forms of visualization, and searching for new sequence pattens based on user-specified features.

## The ScaMot tool

The first fully functional version of the algorithm for large-scale comparative analysis of the effect of point substitutions on motif occurrence has been implemented in ScaMot. The tool was coded in C# based on the .NET platform, so its use was limited to the Windows operating system. It also had limited analytical capabilities, and was practically restricted to generating a binary matrix of motif occurrence and graphical visualizing this matrix. The tool has been used for analysis in several scientific studies, including a published paper ([12], [13], [14]). The executable version of ScaMot (scamot.exe) is available in the PMScanR repository on the GitHub platform in the ScaMot folder.

## Acknowledgements

We would like to thank Mr. Pawel Taborski for his technical support and programming assistance in coding the first version of the algorithm and creating the ScaMot program, which was used during the analyses in an earlier publication. We would also like to thank Professor Stefano Pascarella and Ms. Katarzyna Zyznowska for helping us with the testing of the tool and Professor Jan Sedzik for pointing out the need for an analytical pathway with a key role for protein motifs.

